# Whole genome sequencing reveals the phylogenetic relationship and reticulate evolution of *Phrynocephalus* lizards in Eurasia

**DOI:** 10.1101/2025.09.09.675123

**Authors:** Lin Cui, Roman A. Nazarov, Ying Chen, Yin Qi, Xianguang Guo, Yun Xia, Weizhao Yang

## Abstract

Toad-headed agama (genus *Phrynocephalus*) is a widely distributed group of lizards across Central Eurasia, inhabiting diverse arid and semi-arid environments from the Caspian Basin to the Qinghai–Tibetan Plateau (QTP). Despite considerable research efforts over the past decades, a robust phylogenetic framework for *Phrynocephalus* has remained elusive. In this study, we collected 26 samples representing a substantial portion of the distribution of this genus and established a well-supported phylogeny based on whole-genome sequencing and phylogenomic approaches. We found that the *Phrynocephalus* lizards comprised of three major clades, including the Western Asia, Central Asia and viviparous clades. Furthermore, we provided compelling evidence for an ancient hybridization event that led to the speciation of *P. mystaceus*, which likely contributed to the observed phylogenetic discordance across the genome. Divergence time estimation and ancestral area reconstruction indicated that *Phrynocephalus* lizards originated approximately 31.57 million years ago in the Central Asia and Iranian Plateau region, followed by Miocene dispersals into the Arabian Peninsula, the Gobi Desert, and the QTP. Our results illuminate how hybridization and historical range dynamics have jointly driven the origin and diversification of *Phrynocephalus*, providing new insights into the formation of biodiversity in Eurasia.

## 1. Introduction

Toad-headed agama (Agamidae: *Phrynocephalus* Kaup 1825) is a widespread reptilian taxon across Central Eurasia (Zhao et al., 1999). Its distribution encompasses diverse arid and semi-arid landscapes, including the Arabian Peninsula, Iranian Plateau, Caspian Basin, Turan lowlands, Tarim/Junggar basins, Gobi Desert, and the Qinghai–Tibetan Plateau (QTP) (Macey et al., 2018). These lizards exhibit remarkable adaptive traits to the arid habitats, characterized by dorsoventrally flattened bodies, reduced auditory structures, and specialized scales for burrowing or navigating loose substrates (Arnold, 1999). Since *Phrynocephalus* is both an ecologically important element across Palearctic deserts and a taxon with its diversity still underestimated, studying these lizards can illuminate the processes shaping present-day biodiversity patterns throughout Eurasia.

Although considerable efforts have been made over the past decades, a reliable phylogenetic framework for *Phrynocephalus* lizards has yet to be achieved. In fact, the basal clade in the phylogeny of *Phrynocephalus* remains unresolved. For instance, Solovyeva et al. (2018a) placed the assemblage comprising of *P. arabicus*, *P. longicaudatus*, *P. maculatus* and *P. scutellatusas* as the earliest-diverging lineage within the genus. However, Solovyeva et al. (2023) recovered *P. lutensis* as sister to all remaining species when more samples were added to the analysis. The conflicting results also underscored the controversial origin of the viviparous species in this genus which has not been addressed as well (Macey et al., 2018; Solovyeva et al., 2018a; Solovyeva et al., 2023). In addition, the phylogenetic position of *P. mystaceus* was also under debate, which is a morphologically distinctive species with the largest body size in the genus (Solovyeva et al., 2018b; Whiting et al., 2022). Its singular morphological traits have even led some researchers to treat it at generic rank as *Megalochilus* (von Eichwald, 1831). Early mitochondrial analyses variously placed *P. mystaceus* as sister either to *P. axillaris*, to *P. axillaris* and *P. helioscopus*, or to all other oviparous lineages (Pang et al., 2003). Recent multi-locus analyses documented pronounced mito–nuclear conflict, where *P. mystaceus* was clustered with *P. axillaris* based on mitochondrial DNA (mtDNA), but grouped with other species from Iranian Plateau using nuclear DNA (nuDNA), with AU tests confirming significant incongruence (Solovyeva et al., 2018a). This situation could be partially attributed to the very limited molecular markers used in the phylogenetic studies of *Phrynocephalus* lizards, such as mtDNA or nuDNA fragments. For example, earlier phylogenetic inference relied mostly on a few mtDNA loci (e.g., *COI*/*ND2*/*ND4*/*CYTB*) or short nuDNA fragments (e.g., ∼1,200 bp of *RAG-1*), which often yielded poor resolution (e.g. Pang et al., 2003; Guo and Wang, 2007; Melville et al., 2009; Macey et al., 2018; Solovyeva et al., 2018a; Solovyeva et al., 2023;). In particular, mtDNA and nuDNA often yield statistically significant topological conflicts, as different sources of phylogenetic markers typically represent diverse evolutionary trajectories (Platt et al., 2018). Under these circumstances, whole genome-wide data was required to reveal the phylogenetic framework of *Phrynocephalus* lizards.

Hybridization is another major source of phylogenetic conflicts (Steenwyk et al., 2023). In fact, hybridization has been found to be prevalent in nature, with at least 25% of the plants and 10% of the animals having experienced hybridization (Mallet, 2005; Runemark et al., 2019). This phenomenon resulted in “mosaic genomes” of species, where different genomic fragments may have undergone different histories, hindering the phylogenetic reconstruction when only a few molecular markers were used (Steenwyk et al., 2023). For example, heliconian butterflies (Edelman et al., 2019), swordtail fishes (Schumer et al., 2018), gray snub-nosed monkey (Wu et al., 2023a), and cultivated potato (Zhang et al., 2025) all showed evidence of extensive hybridization, and the phylogenetic networks may better accommodate these reticulate signals than bifurcating trees (Huson and Bryant, 2006). Evidence of hybridization has also been detected in *Phrynocephalus* lizards. Gao et al. (2022) identified recent asymmetric introgression from P*. putjatai* into *P. vlangalii* at the Gonghe contact zone. In addition, Guo et al. (2025) further inferred multiple ancient reticulation events within the viviparous clade, including around 41% of alleles in *P. erythrurus* from the western lineage of *P. vlangalii* and 14% in *P. theobaldi* from *P. erythrurus*. However, the pattern of ancient hybridization has not been systematically assessed throughout the *Phrynocephalus* genus, limiting further understanding of the evolutionary history for this group of lizards.

In this study, we conducted an integrative investigation to illustrate the evolutionary history of *Phrynocephalus* lizards using whole genome sequencing data. We have three main objectives: (1) to reconstruct a robust phylogenomic framework for *Phrynocephalus* lizards; (2) to estimate the ancient reticulate events between the species; and (3) to reveal the origin and diversification process of *Phrynocephalus* species. To achieve these aims, we collected samples from a considerable portion of the distribution of those species, including Arabian Peninsula, Iranian Plateau, Central Asia, Tarim/Junggar Basin, Gobi Desert and QTP. Through whole genome sequencing and phylogenomic analysis, we present a robust phylogenetic framework for *Phrynocephalus* and provide strong evidence linking its phylogenetic discordance of major clades to ancient hybridization, thereby revealing how hybridization and historical range dynamics facilitated the origination and diversification of biodiversity across arid habitats in Eurasia.

## 2. Materials and Methods

### 2.1. Animal Ethics

Animal collection procedures followed all relevant laws and regulations, as well as the applicable guidelines for animal care and use. Sampling and experiments were approved by the Ethical Committee for the Animal Experiments in Chengdu Institute of Biology, Chinese Academy of Sciences, with Permission No. CIBDWLL2022025. *Phrynocephalus* samples from Iran, Jordan, Oman, Uzbekistan and Kazakhstan were provided by the Zoological Museum of Moscow State University (ZMMU), with corresponding permission for genomic research granted by the administration and the collection curator of ZMMU (Permission No. ZM-027).

### 2.2. Sampling and Sequencing

A total of 26 samples were used in this study, representing 22 *Phrynocephalus* species and two outgroup species: *Pogona vitticeps* and *Laudakia wui*. Among them, 20 samples were newly collected: 19 *Phrynocephalus* samples encompassing one subspecies of *P. helioscopus* (*P. h. sergeevi*) and one newly described subspecies of *P. mystaceus* (*P. m. khorasanus*; Solovyeva et al., 2018b), as well as one *L. wui* sample. All sample details are provided in Table S1. Genomic DNA was isolated from toe tissues cryopreserved at -80°C using DNeasy Blood and Tissue Kit (Qiagen, Valencia, CA, USA). Short-read sequencing was performed on Illumina Novaseq X platform (Novogene, Beijing, China) generating 150 bp paired-end reads. All raw sequencing reads have been deposited in the National Genomics Data Center (BioProject accession: PRJCA045046). In addition, sequence reads of five *Phrynocephalus* species were downloaded from National Genomics Data Center (accession No. PRJCA022059), and one sample of *P. vitticeps* was downloaded from European Nucleotide Archive (accession No. ERZ094017).

### 2.3. Reference Genome Optimizing

To minimize the reference bias in downstream variants calling, we generated a reference genome comprising only syntenic segments shared by four species with published high-quality genomes, including *P. vlangalii* (NGDC accession No. PRJCA022059), *P. guinanensis* (NGDC accession No. PRJCA027608), *P. versicolor* (NCBI accession No. PRJNA738137) and *P. vitticeps* (ENA accession No. ERZ094017). Pairwise whole-genome alignments were first conducted between *P. vlangalii* and each of the other three assemblies using LASTZ (v1.04.00; Harris, 2007) with the following parameters: --step=19, --hspthresh=2200, --inner=2000, --ydrop=3400, -- gappedthresh=10000, --format=axt. High-scoring AXT blocks were then chained and netted with the UCSC tools axtChain and chainNet (-minScore=5000; -linearGap=loose) to obtain one-to-one, non-overlapping orthologous regions (Hinrichs et al., 2006). The resulting pairwise nets were merged with MULTIZ (v11.2; Blanchette et al., 2004) to obtain a four-way multiple alignment. Only blocks present in all four genomes were retained, whereas lineage-specific or non-syntenic regions were soft-masked to “N”, yielding a masked genome that served as the mapping reference for variant discovery.

### 2.4. Variants Calling

Raw reads were subjected to quality trimming and adapter clipping by Trimmomatic with default settings (v0.39; Bolger et al., 2014). The processed sequence reads were aligned to the reference genome using BWA (v0.7.17; Li, 2013). PCR duplicates were subsequently marked by the “Mark Duplicates Spark” module in GATK (v4.5.0.0; Van der Auwera and O’Connor, 2020). Single nucleotide variants (SNVs) calling was performed according to GATK best practices pipeline, followed by variant hard-filtering with the following parameters: QD < 2.0, MQ < 40.0, FS > 60.0, SOR > 3.0, MQRankSum < -12.5, ReadPosRankSum < -8.0 with GATK “Variant Filtration” module. Sites within 5 bp of indels and clustered variants (inter-site distance below 5 bp) were also filtered by GATK “Variant Filtration” with flags: --mask-extension=5, --cluster-window-size=5, --cluster-size=2. Additional filtering was implemented through VCFtools (v0.1.17; Danecek et al., 2011), including removing variants with mean depth < 8× and sites with depth greater than twice the mean depth to eliminate low-depth or potential paralogous variants.

### 2.5. Phylogenomic Analysis

We employed two strategies, concatenation and multispecies coalescent (MSC), to infer a robust phylogenomic framework for *Phrynocephalus* lizards. *P. vitticeps* and *L. wui* were used as outgroup. All phylogenetic relationships were inferred using IQ-TREE (v1.6.12; Nguyen et al., 2015) under the GTR+ASC model with 1,000 bootstrap replicates. For the concatenation approach, we generated two concatenated SNV alignments: one comprising all SNVs and the second comprising SNVs located in protein-coding regions, yielding the WGS-Tree and the CDS-Tree, respectively. For the coalescent approach, we partitioned the genome into fixed-length windows with three different sizes: 50kb, 100kb and 200kb. Local trees for each window were first inferred, and filtered local trees with mean bootstrap values below the thresholds of 80. The resulting pruned trees were then subjected to ASTRAL-III (v5.7.1; Zhang et al., 2018) to estimate the MSC trees. To quantify topological incongruence among trees generated by the different phylogenetic analyses, we computed normalized Robinson–Foulds distances (Robinson and Foulds, 1981) for every pairwise comparisons with the R package “ape” (v5.8-1; Paradis and Schliep, 2019). To further illustrate the phylogenetic discordance across the genome, we utilized TWISST (Martin and Van Belleghem, 2017) to check the topology weighting of local trees in 200kb windows. Python script “twisst.py” and R script “plot_twisst.R” (https://github.com/simonhmartin/twisst) were used to calculate and plot the results from. In addition, a DensiTree plot was generated using DensiTree (v3.1.0; Bouckaert and Heled, 2014).

Moreover, mitochondrial genomes of 26 agamid species were assembled by MEANGS (v1.2.1; Song et al., 2022) from raw sequence reads with default parameters and annotated by MitoFinder (v1.4.2; Allio et al., 2020). Then, 12 protein coding genes (*ATP8* was excluded due to annotation failure in several samples) were subsequently aligned, trimmed, and concatenated using PhyloSuite (v1.2.3; Zhang et al., 2020), followed by removal of third codon positions in case of substitution saturation. Thereafter, the best-fitting nucleotide substitution model was selected by ModelFinder (v2.2.0; Kalyaanamoorthy et al., 2017) and subsequently implemented in IQ-TREE (v1.6.12; Nguyen et al., 2015) to infer the maximum likelihood Mito-Tree reconstruction.

### 2.6. Hybridization Detection

We applied three approaches to detect gene flow between *Phrynocephalus* species. We first utilized the “Dtrios” module of Dsuite (v0.5r58; Malinsky et al., 2021) to compute Patterson’s D statistic (ABBA-BABA test; Patterson et al., 2012) for all possible trios of species and their inferred ancestors. Subsequently, using Dsuite’s “Fbranch” module (v0.5r58; Malinsky et al., 2021), we conducted an *f-*branch analysis based on the D statistic to evaluate gene flow across the phylogeny. Because viviparous species exhibit extensive hybridization (Guo et al., 2025), we retained only a single viviparous species to sharpen signal detection among the remaining taxa.

Following, we applied TreeMix (v1.1; Pickrell and Pritchard, 2012) to infer both the strength and direction of gene flow. Specifically, genome-wide allele frequency data were generated using PLINK (v1.90b7.2; Chang et al., 2015) and converted into a format compatible with TreeMix via the “plink2treemix.py” script available at https://bitbucket.org/nygcresearch/treemix/downloads/. Then, a maximum likelihood populations tree was generated by TreeMix, which demonstrated an identical topology to WGS-Tree. Subsequently, in-house scripts were used to conduct five replicate analyses for each of the species under migration scenarios ranging from 1 to 10 based on the tree above. The optimal migration scenario was determined using the R package “OptM” (v0.1.8; Fitak, 2021) (https://github.com/cran/OptM) with both “Evanno” and “Linear” methods. Finally, the TreeMix plot was generated under this single-migration scenario with the “plotting_funcs.R” script (https://bitbucket.org/nygcresearch/treemix/src/master/src/).

We also utilized ADMIXTOOLS 2 (v2.0; Patterson et al., 2012; Maier et al., 2023) in R to reconstruct the admixture graph to visualize the reticulate evolutionary history and quantify parental contributions to hybrids. The “find_graphs” function was used to automatically identify the best-fitting graph under a single admixture event, based on a set of *f*3-statistics derived from precomputed *f*2-statistics. In addition, the “qpGraph” function was used to compute admixture scores for three alternative models designed based on phylogenetic relationship without hybridization for comparative validation.

### 2.7. Divergence Dating and Ancestral Area Reconstruction

To reduce bias arising from historical introgression in node-age estimation, we confined molecular dating to genomic segments with local topologies concordant with the WGS-Tree. This filtering yielded 1,003,564 SNVs. Due to the scarcity of fossil records for *Phrynocephalus* lizards, two soft constraints (C1 and C2) were applied according to the two well-dated paleogeographic events: C1 targeted the crown of the viviparous clade and constrained its origin within a conservative time window aligned with the establishment of the Taklimakan Desert, implemented as a soft maximum at ∼7.0 Ma (Sun et al., 2009); C2 constrained the split between *P. helioscopus* and the remaining oviparous taxa with an upper bound near 11 Ma, consistent with the onset of major Tian Shan uplift (∼10–11 Ma; Charreau et al., 2009). Divergence times were estimated with “MCMCTree” program in the PAML package (v4.9j; Yang, 2007) under the independent-rates relaxed-clock model. Each analysis was run for 30,000,000 iterations with sampling every 2,000 steps and treated former 50% iterations as burn-in. The MCMC analysis showed good convergence. Posterior means and 95% highest posterior density (HPD) intervals for node ages were extracted from “MCMCTree” outputs.

We inferred ancestral distribution and biogeographic processes shaping the present-day pattern of *Phrynocephalus* using BioGeoBEARS (v1.1.1; Matzke, 2013) on the time-calibrated phylogeny. Six alternative models were compared: DEC (Dispersal–Extinction–Cladogenesis), DIVALIKE (likelihood version of Dispersal–Vicariance Analysis), and BAYAREALIKE (likelihood version of Bayesian Inference of historical biogeography for discrete areas), together with their +J counterparts that incorporate founder-event speciation (Matzke, 2014). Model fit was assessed with AICc, and the best-fitting model and parameter estimates were reported (dispersal *d*, extinction *e*, and founder event *j* when applicable). For geographic coding, we adopted a five-area scheme based on Solovyeva et al. (2018a): AP (Arabian Peninsula), CA (Central Asia), IP (Iranian Plateau), QTP (Qinghai–Tibetan Plateau), and EGD (Extensive Gobi Desert). We limited the maximum range size to two areas per lineage to prevent unrealistically widespread ancestral ranges.

**Figure 1.**
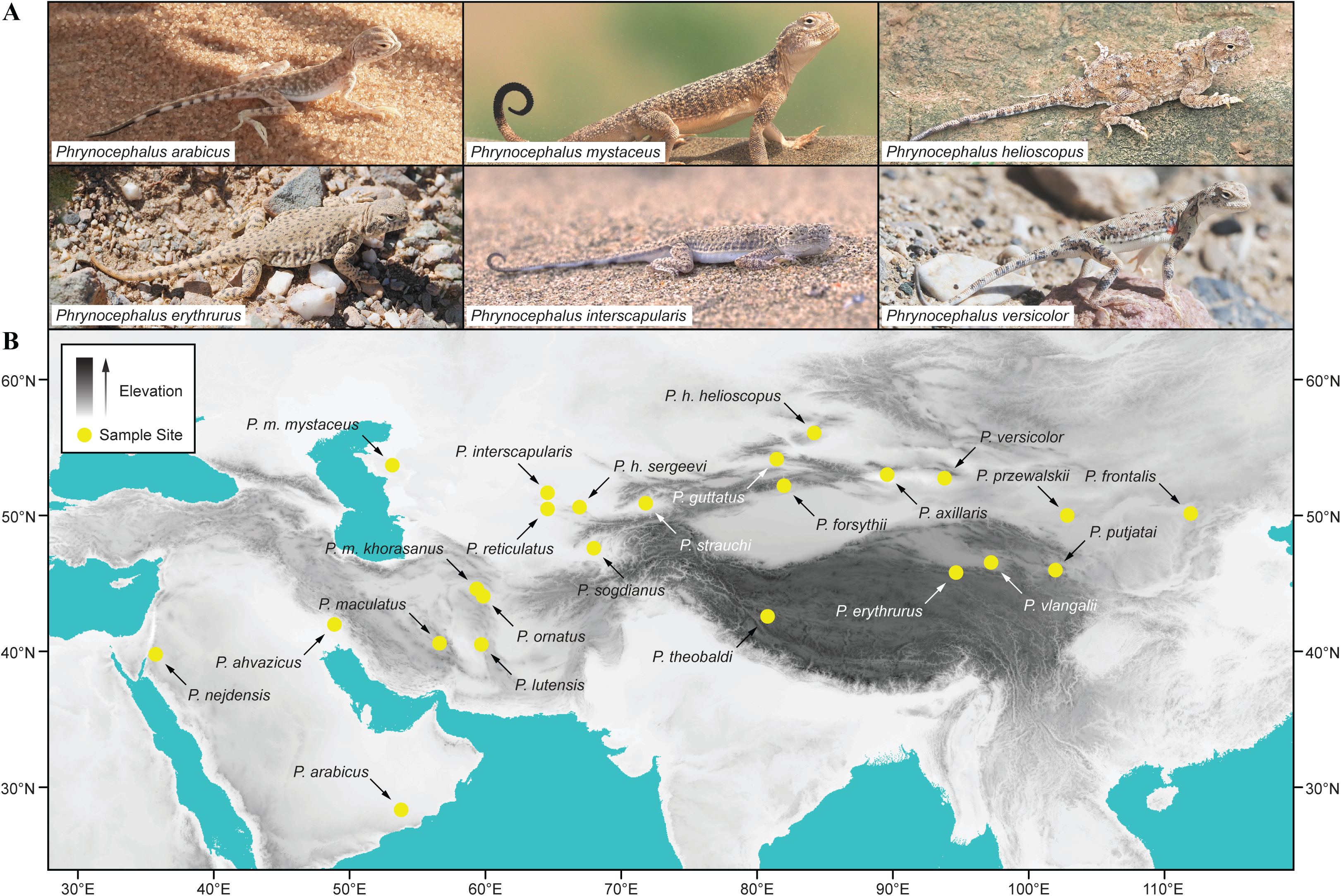
Diverse phenotypes and sampling sites of *Phrynocephalus*. (A) Photographs of representative *Phrynocephalus*, and (B) geographic sampling localities of the individuals analyzed in this study.

## 3. Results

### 3.1. Variants Calling

A total of 26 agamid genomic data were mapped to the optimized reference genome, achieving an average mapping rate of 67.4%. After stringent filtering, a total of 4,889,121 high-quality SNVs with a mean depth of 14.76× per site were obtained.

### 3.2. Phylogenetic Framework

For concatenation approach, both WGS-Tree (Figure 2B) and CDS-Tree (Figure S1) successfully reconstructed three major clades within *Phrynocephalus* lizards: the Western Asia (WA) clade, the Central Asia (CA) clade, and the viviparous (VP) clade. The two maximum likelihood trees exhibited identical inter- and intra-clade topologies, with all bootstrap values reaching 100% except for the node uniting *P. erythrurus* and *P. theobaldi* within the VP clade in the CDS-Tree (bootstrap value = 53%; Figure S1). Under MSC approach, all three window-based MSC-Trees (50kb, 100kb, and 200kb) also recovered the same three major clades of WA, CA, and VP (Figure S2). The 50kb and 100kb coalescent trees were topologically identical to the concatenation trees, as confirmed by a Robinson–Foulds pairwise tree distance of 0 to both the WGS-Tree and the CDS-Tree (Figure 2A). In contrast, the 200kb MSC-Trees differed only in the placement of the deepest split: 200kb MSC-Trees supported the WA clade as sister to the CA clade, whereas the 50kb and 100kb MSC-Trees and both concatenation trees favored the VP clade as the sister group to the CA clade (Figure 2B). Apart from this single conflict, internal relationships within each major clade were congruent across approaches and window sizes.

**Figure 2.**
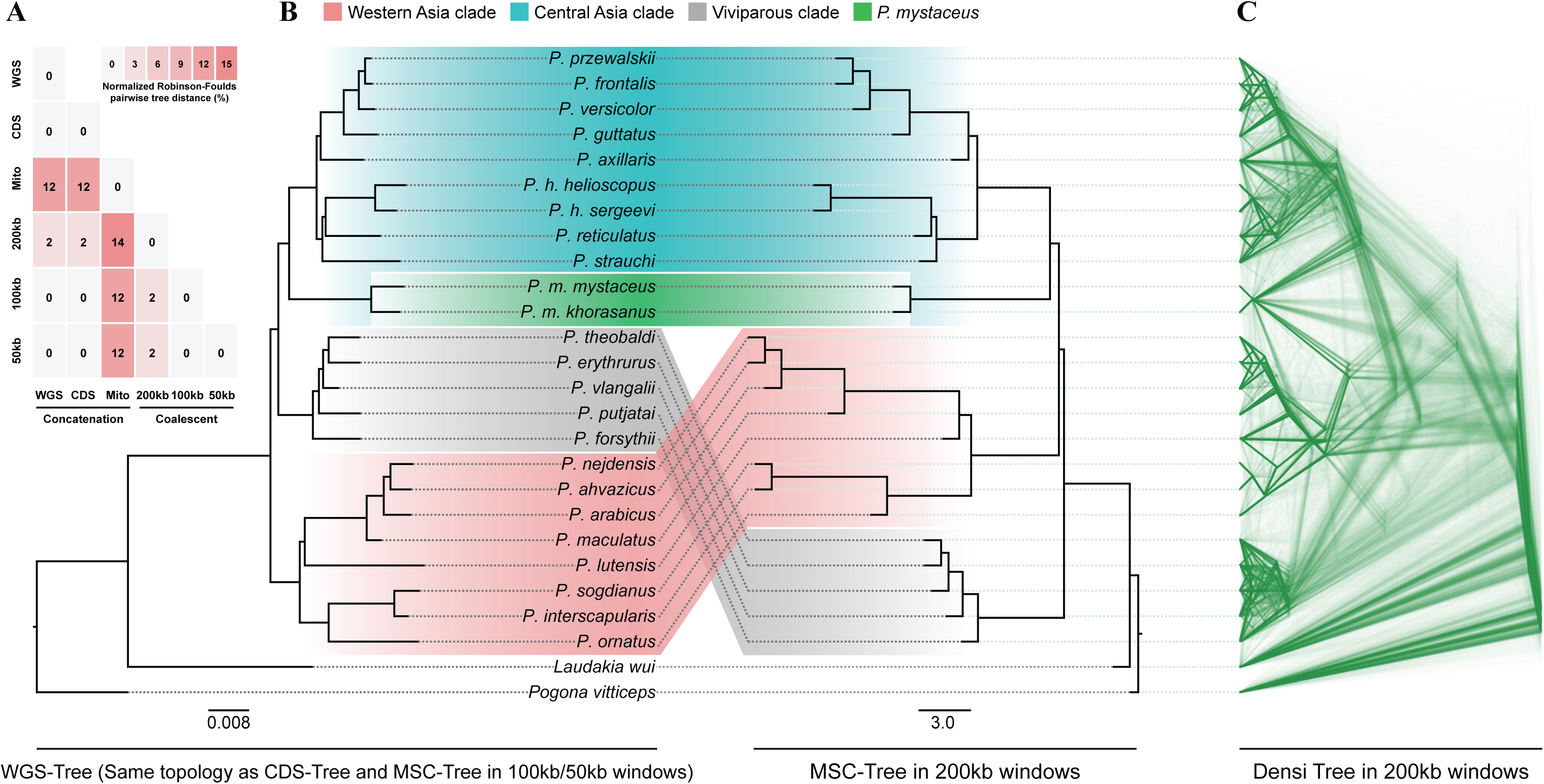
Phylogenetic discordance between coalescent and concatenation approaches. (A) A summary of the normalized Robinson–Foulds distances between pairwise comparisons of phylogenies. (B) The topological discordance between the concatenation tree and the coalescent tree, where all nodes demonstrate a bootstrap support value of 100%. (C) DensiTree in an alignment of 8,662 local tree of 200kb windows, illustrating genome-wide phylogenetic discordance.

DensiTree visualization for the 200kb windows revealed extensive heterogeneity among local topologies (Figure 2C). Further topological analysis of local trees recovered fifteen distinct inter-clade topologies among the three major clades and the *P. mystaceus* (Figure S3A); the three most prevalent topologies were topology1 (20.14%), topology2 (17.83%) and topology4 (13.35%) (Figure S3B), corresponding respectively to the coalescent topology, the concatenation topology, and a topology in which *P. mystaceus* was sister to the WA clade (whereas in topology1 and topology2, *P. mystaceus* was sister to the CA clade). Based on this, we excluded the VP clade and focused on the remaining two major clades and *P. mystaceus* (Figure 3B). Proportional analysis revealed that more than half (56%) of the local trees supported the *P. mystaceus* was sister to the CA clade (Figure 3C); 33% supported the *P. mystaceus*–WA clade pairing; and 11% supported the CA clade–WA clade pairing. Moreover, across the genome these three topologies were evenly distributed, with no genomic region showing preferential enrichment of any particular topology (Figure 3A).

**Figure 3.**
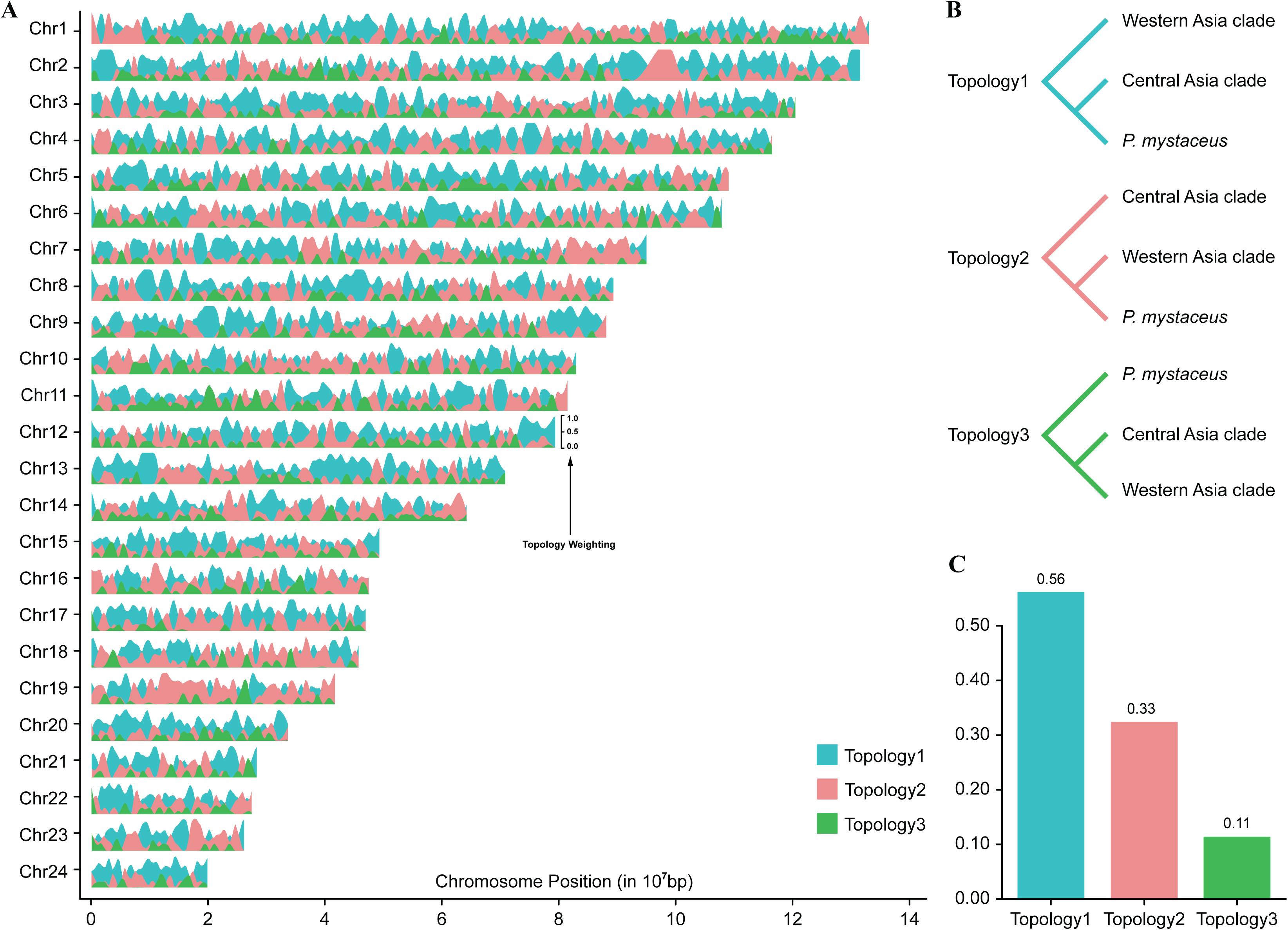
Chromosome-scale topology weighting reveals pervasive phylogenetic discordance. (A) Smoothed TWISST weightings for the focal topology across the 24 chromosomes. (B) Schematic of the three alternative topologies relating the two major clades and the *P. mystaceus*. (C) Genome-wide distribution of the relative weightings assigned to each topology.

Mito-Tree (Figure S4) exhibited topological discordance with WGS-Tree, with normalized Robinson–Foulds pairwise tree distances ranging from 12% to 14% (Figure 2A). Specifically, we found that in both the WGS-Tree and the CDS-Tree, *P. mystaceus* occupied the basal position within the CA clade with strong support, whereas in the Mito-Tree the basal position of the CA clade was held by *P. axillaris* (bootstrap value = 90%); and in the WA clade of the Mito-Tree, *P. ahvazicus* was inferred to be sister to *P. arabicus* (bootstrap value = 61%), whereas the nuclear trees supported a *P. ahvazicus*–*P. nejdensis* pairing.

### 3.3. Genomic Evidence for Ancient Hybridization

We first applied Patterson’s D-statistic to all 2,024 triplets with *L. wui* as the outgroup. A total of 79.30% of triplets showed significant deviation from the null of no introgression (Z score > 3), indicating that interspecific gene flow, rather than incomplete lineage sorting, underlay the observed phylogenetic discordance. Furthermore, the *f*-branch results showed that three particularly strong signals were evident: (1) the ancestral branches of both *P. mystaceus* induvials showed strong gene flow with all WA-clade species; (2) a pronounced introgression signal between *P. reticulatus* and *P. strauchi*; and (3) the strong gene flow signal between *P. vlangalii* and the ancestor of the CA clade.

In TreeMix analysis, both the “Evanno” and “Linear” criteria supported migration edge = 1 as the optimal scenario (Figure S5). In this scenario, strong gene flow was detected from the most recent common ancestor (MRCA) of the WA clade into *P. mystaceus* (migration edge weight = 0.45) (Figure 4B), indicating that approximately 45% of *P. mystaceus* alleles were introgressed from the WA clade.

**Figure 4.**
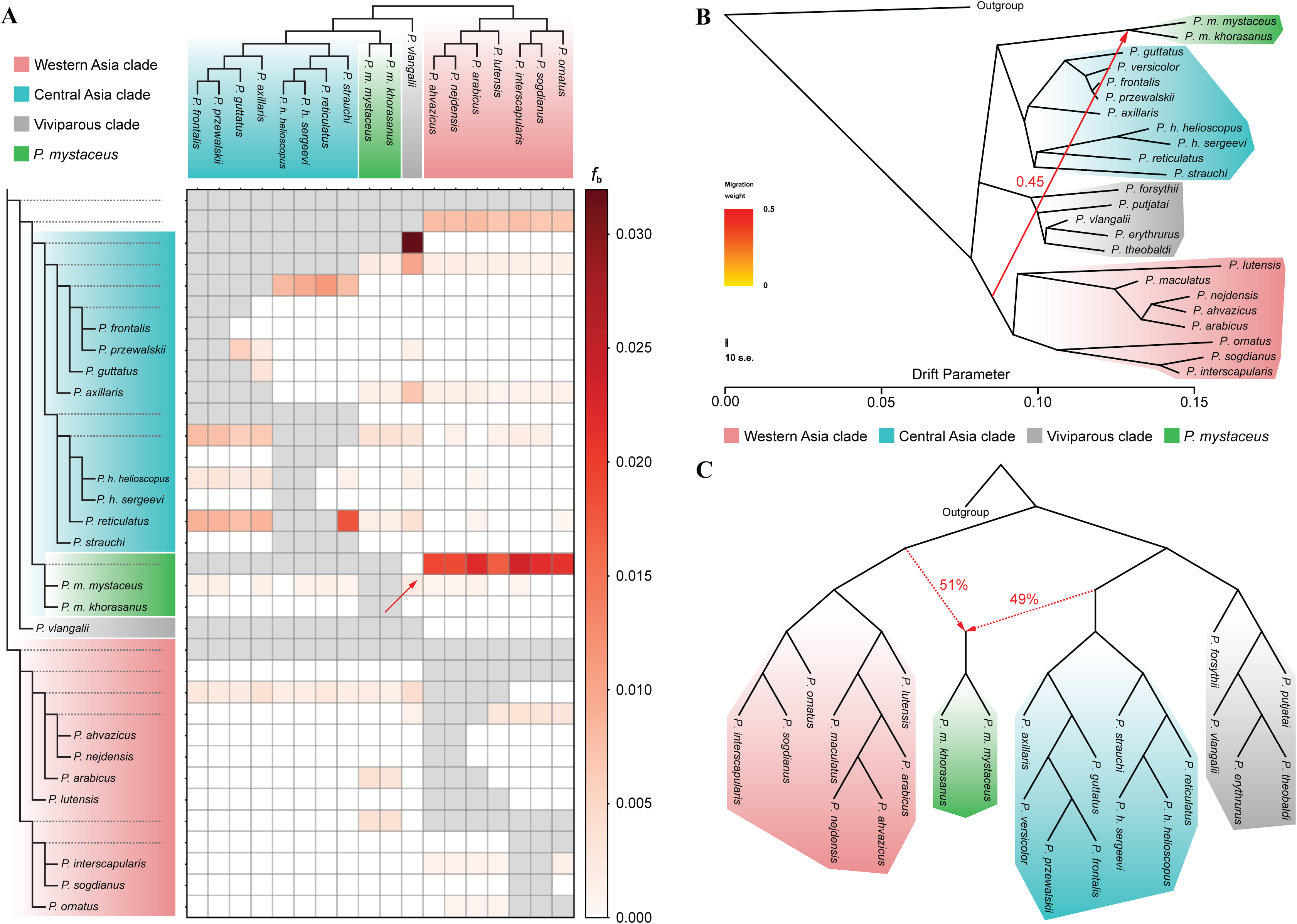
Evidence for an ancient hybrid origin of *P. mystaceus*. (A) *f*-branch analysis summarizing significant interspecific gene flow among 18 extant taxa and their reconstructed ancestors. Dashed lines denote ancestral nodes, and the red arrow marks the ancestral node leading to both *P. mystaceus* individuals, which exhibits pronounced gene flow with the Western Asia clade. (B) TreeMix maximum likelihood phylogeny with one migration edge (standard-error bars provided) shows excess allele sharing from the Western Asia clade to *P. mystaceus* (migration edge weight = 0.45; red arrow). (C) Admixture graph inferred with ADMIXTOOLS 2 supports an ancient hybrid origin of *P. mystaceus*, with nearly equal ancestry contributions from the ancestors of the Western Asia and Central Asia clade (51% vs. 49%).

The “find_graphs” function in ADMIXTOOLS 2 identified an admixture graph under one hybridization event with a score of 4222.41 (Figure 4C, also see Model 1 in Figure S6). Furthermore, three additional no-hybridization models as provided in Figure S6 were evaluated using the “qpGraph” function: (1) *P. mystaceus* as sister to either putative ancestor (Model 2 with a score of 9458.97 and Model 3 with a score of 6104.60), (2) MSC-Tree topology (Model 3), and (3) WGS-Tree topology (Model 4 with a score of 9855.85), demonstrated that the lowest-scoring single-admixture model was superior. Consistent with the TreeMix result, these results indicate that *P. mystaceus* originated from ancient hybridization between the WA and CA clade ancestors, with approximately equal ancestry (51% from WA; 49% from CA).

### 3.4. Divergence Dating and Ancestral Distribution

The time-calibrated phylogeny (Figure 5A) placed the origin of *Phrynocephalus* between the late Eocene and early Miocene, with a median estimate of 31.57 Ma (95% HPD: 20.23–43.51 Ma), and dated the first divergence to form the CA and IP clades in the early–middle Miocene at approximately 17.63 Ma (95% HPD: 12.62–22.65 Ma), then the second split to form the VP clade dated into the middle–Miocene at approximately 15.81Ma (95% HPD: 11.39–20.47 Ma). Other divergence times for key nodes can be found in Table S2.

**Figure 5.**
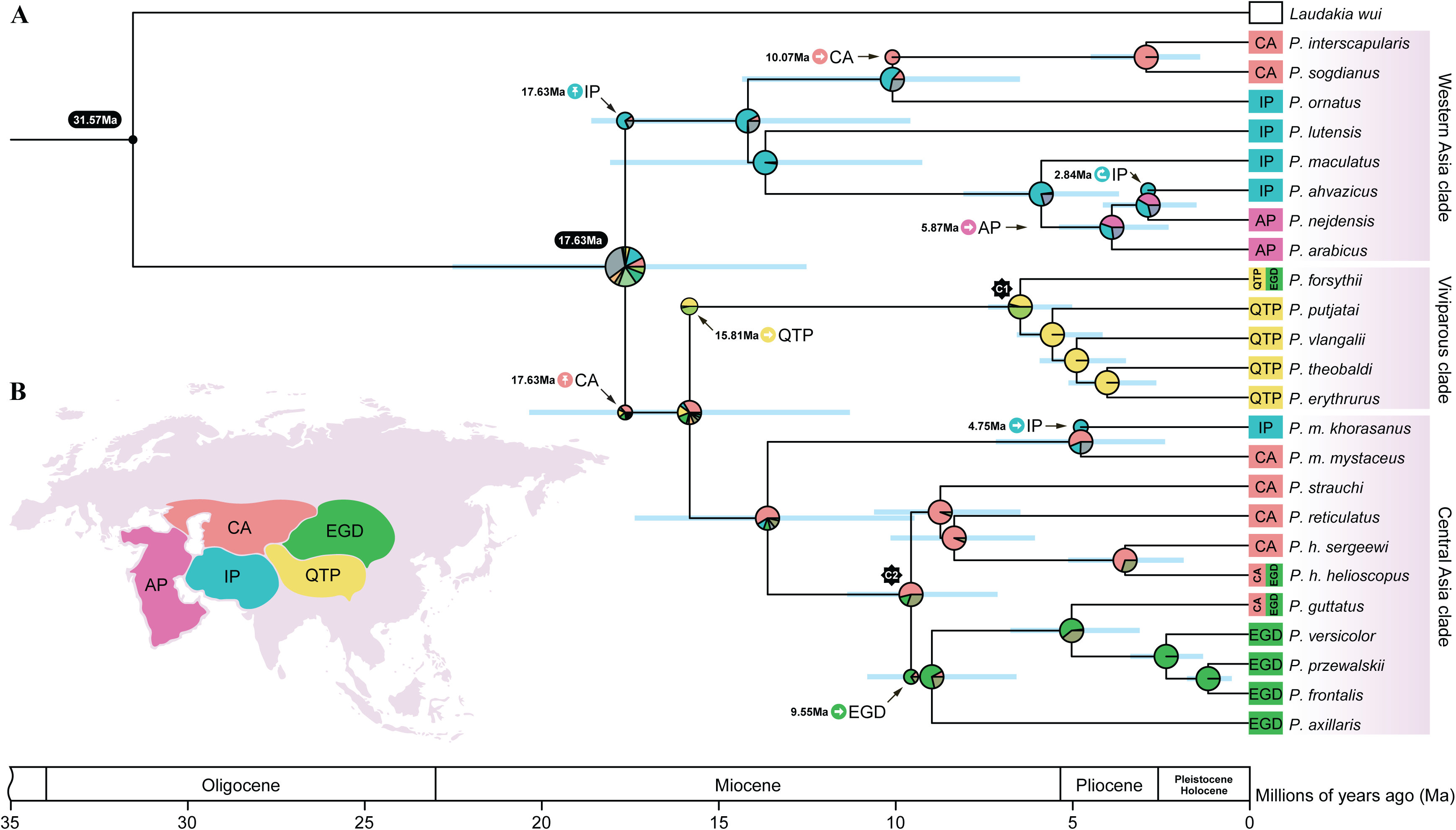
Spatiotemporal reconstruction of *Phrynocephalus*. (A) Divergence time estimation with two soft calibrate point (asterisk C1 and C2), and ancestral area reconstruction based on the best-fitting model (DEC + J). Abbreviations: AP, Arabian Peninsula; CA, Central Asia; IP, Iranian Plateau; QTP, Qinghai–Tibetan Plateau; EGD, Extensive Gobi Desert. (B) Illustration of biogeographic areas used in this study.

In ancestral area reconstructions, DEC+J was identified as the best fitting model (AICc = 72.52; Table S3). Under this model, the MRCA of *Phrynocephalus* was inferred to have occupied Central Asia (CA) together with the Iranian Plateau (IP). From this source area, the first divergence to form three major clades (the CA, WA, and VP clade) started during the Miocene. The reconstructions further indicate a back dispersal of *P. guttatus* from EGD (Extensive Gobi Desert) to CA near the onset of the Pliocene (Figure 5).

## 4. Discussion

Toad-headed agama (genus *Phrynocephalus*) is a symbolic reptilian taxon in central Eurasia, distributing in the arid and semi-arid habitats from the Caspian Basin to East Asia. However, a robust phylogenetic framework remains largely unresolved, which hinders further understanding of the origin and diversification of this genus. In this study, we collected 26 samples from a considerable portion of the distribution area of *Phrynocephalus* lizards. By conducting whole genome sequencing and integrated phylogenomic analysis, we reconstructed the evolutionary history of *Phrynocephalus* lizards, with an ancient reticulate event which led to the hybrid origin of *P. mystaceus*. Through divergence time and ancestral states estimation, we also inferred that the *Phrynocephalus* lizards might have originated from Central Asia and Iranian Plateau at approximately 31.57 Ma and then split into three major clades that subsequently spread to other regions.

Our study implemented whole genome sequencing data and phylogenomic approaches to reconstruct the evolutionary history of *Phrynocephalus* lizards, which generated the phylogenetic framework between species. Currently, genome-wide data have been commonly used to infer the phylogenetic relationship for species that were difficult to resolve using only the short markers. For example, the long-debated unresolved relationships between flightless ratites and volant tinamous (Cloutier et al., 2019), mbuna cichlids (Scherz et al., 2022), and manakins (Zhao et al., 2023) have been addressed based on genome-wide markers. In this study, we performed both the concatenation and multispecies coalescent strategies based on more than 4.88 million variants, obtaining a robust phylogenetic framework for this genus. Our results clearly showed that the *Phrynocephalus* species can be divided into three major clades: the WA (Western Asia) clade, the viviparous clade, and the CA (Central Asia) clade. We also found that the viviparous specie was split from the oviparous species, which indicated that the oviparity was the ancestral trait of *Phrynocephalus* lizards. Our results were quite different from previous views, in which earlier broad-scale analyses recovered nine high-support species groups (Solovyeva et al., 2014); an updated synthesis integrating mtDNA, nuDNA, and allozyme data recovered 13 well-supported *Phrynocephalus* clades (Macey et al., 2018); subsequent mtDNA analyses recognized ten major species groups; and the most recent study supported the monophyly of six species groups (Solovyeva et al., 2023). The discrepancy probably resulted from the unresolved phylogenies from previous studies inferred based on only a few fragments, such as mitochondrial genes or nuclear fragments.

Our results provided strong evidence of an ancient reticulate event where the *P. mystaceus* originated from hybridization between the ancestors of the WA and CA clades with almost equal contribution. In fact, along with the increasing number of cases being detected, hybridization has been considered prevalent in nature (Runemark et al., 2019), and intensifying human activities may be driving additional hybridization events (Wei et al., 2024). Accordingly, phylogenetic reconstruction may be difficult for species that have experienced hybridization. For instance, genome-scale analyses of *Heliconius* butterflies showed that rampant introgression produced pervasive gene tree discordance and undermined support for a single bifurcating species tree, thereby obscuring deep relationships (Edelman et al., 2019). Likewise, genus-wide phylogenomics of *Xiphophorus* swordtails showed that hybridization preceded speciation and generated mito-nuclear discordance, affecting species-tree reconstruction (Du et al., 2024). In our case, historical gene flow involving *P. mystaceus* likely drove discordance in species tree inference, manifested as a genome-wide mosaic of three alternative loca-tree topologies. The hybrid origin of *P. mystaceus* may have also complicated reconstruction of its phylogenetic relationship with other species. For example, Pang et al. (2003) generated the first comprehensive mtDNA phylogeny for Chinese *Phrynocephalus* and observed conflicting topologies that *P. mystaceus* was variably positioned as sister to *P. axillaris*, basal to *P. axillaris* and *P. helioscopus*, or even as the most basal oviparous lineage, depending on whether maximum parsimony, neighbor joining, maximum likelihood or Bayesian inference was performed. Then, Guo and Wang, (2007) revealed that *P. mystaceus* formed a strongly supported basal position relative to all other Chinese species, highlighting its deep divergence and aligning with earlier morphological hypotheses that treated *P. mystaceus* as a distinct genus (von Eichwald, 1831). In this study, we implemented multiple approaches to estimate the reticulate events, including D statistic, *f*-branch, TreeMix and qpGraph. Our results strongly supported the origin of *P. mystaceus* through hybridization between the ancestors of the WA clade (51% of ancestry contribution) and CA clade (49% of the ancestry contribution). Actually, Solovyeva et al. (2018a) identified the mito-nuclear discordance of *P. mystaceus* and interpreted it as potential evidence of ancient interspecific hybridization. Therefore, our study supported that a “web of life” thought could be more appropriate than a “tree of life” to reconstruct the evolutionary history of species.

Our results showed that *Phrynocephalus* lizards diverged from other Agamid species at approximately 31.57 Ma (95% HPD: 20.23–43.51 Ma), with CA+IP inferred as the most probable ancestral area. A basal radiation followed in the early–middle Miocene (ca. 17.63 Ma; 95% HPD: 12.62–22.65 Ma), with subsequent range expansions from CA to the QTP in the middle Miocene (ca. 15.81 Ma; 95% HPD: 11.39–20.47 Ma), and later spread to the EGD (ca. 9.55 Ma; 95% HPD: 7.20–11.46 Ma) and also the AP (ca. 5.81 Ma; 95% HPD: 3.76–8.17 Ma). Despite the largest arid zone in the world, Central Asia and Irano-Anatolian are indeed a biodiversity hotspot of the Eurasian continent (Noroozi et al., 2021; Vintsek et al., 2024). Recent floristic syntheses indicate that Central Asia sustains high vascular plant richness with substantial endemic fractions across national floras, corroborating its status as a regional biodiversity hotspot (Ma et al., 2024). Fine-scale endemicity analyses in Irano-Anatolian reveal “hotspots-within-hotspots” concentrated in high mountain ranges (e.g., the Alborz and Zagros mountains), with endemic richness and endemicity increasing with topographic complexity and elevation (Noroozi et al., 2018). However, biogeographic studies are still lacking to document how species evolved in this region. Only a few species, such as notably Iranian snakes (Moradi et al., 2024) and *Mesalina watsonana* species complex in the Iranian plateau (Boroumand et al., 2024), as well as widespread *Eremias velox* (Liu et al., 2019) and *P. helioscopus* in arid Central Asia (Wu et al., 2023b), yet most studies remain regional in scope. As a symbolic taxon of deserts, our results showed that the *Phrynocephalus* lizards first diversified in CA+IP, along with along with the Middle Miocene Climatic Transition (MMCT; ca. 13.8–14.7 Ma). Across the MMCT, global cooling and steeper latitudinal temperature gradients coincided with a reorganization of atmospheric circulation that promoted steppe–desert expansion and progressive aridification in the Eurasian interior, especially Central Asia (Steinthorsdottir et al., 2021). In this climatic context, increasing aridity and expansion of sand substrates likely catalyzed the initial split of *Phrynocephalus*. Then, *Phrynocephalus* spread to colonize the QTP during ca. 15.81 Ma, a period that coincided with hypothesized major uplift phases of the QTP during the mid-to late Miocene (Miao et al., 2022). Our study not only provided a framework to illustrate how *Phrynocephalus* lizards evolved out of Central Asia and Iranian Plateau to colonize the broad arid habitats in Eurasia but also offered novel insights into the understanding of how biodiversity formed in this region.

## 5. Conclusion

In this study, combined with whole-genome sequencing and phylogenomic approaches, we reconstructed a robust phylogenetic framework for *Phrynocephalus* lizards, which comprised of three major clades, including Western Asia, Central Asia and viviparous clades. In addition, we provided strong evidence for an ancient hybridization event that led to the speciation of *P. mystaceus*. Divergence time estimation and ancestral area reconstruction indicated that this genus originated approximately 31.57 million years ago in the Central Asia and Iranian Plateau region, followed by Miocene dispersals into the Arabian Peninsula, the Gobi Desert, and the QTP. Our results demonstrated how hybridization and historical range dynamics have jointly driven the origin and diversification of *Phrynocephalus*, providing new insights into the formation of biodiversity in Eurasia.

## Supporting information

Spplementary Materials

## CRediT Authorship Contribution Statement

**Lin Cui:** Writing – original draft, Methodology, Investigation, Formal analysis. **Roman A. Nazarov:** Writing – review & editing, Investigation. **Ying Chen:** Writing – review & editing, Investigation, Formal analysis. **Yin Qi:** Writing – review & editing, Investigation. **Xianguang Guo:** Writing – review & editing, Investigation. **Yun Xia:** Writing – review & editing, Investigation. **Weizhao Yang:** Writing – review & editing, Funding acquisition, Data curation, Conceptualization.

## Acknowledgements

We thank Z. Yao for assistance with photographs of *Phrynocephalus*. We are grateful to Dr. Y. Zheng, D. Kang, Y. Wang, Q. Xu, and Y. Tian for constructive comments and suggestions on earlier drafts of this manuscript.

## Data Availability

All sequencing data have been deposited in the Genome Sequence Archive (GSA) of the National Genomics Data Center under Accession number PRJCA045046.

