## Supplementary material for "Whole genome sequencing reveals the phylogenetic relationship and reticulate evolution of *Phrynocephalus* lizards in Eurasia": Spplementary Materials

This Supplementary information contains:

6 Supplementary Figures

3 Supplementary Tables

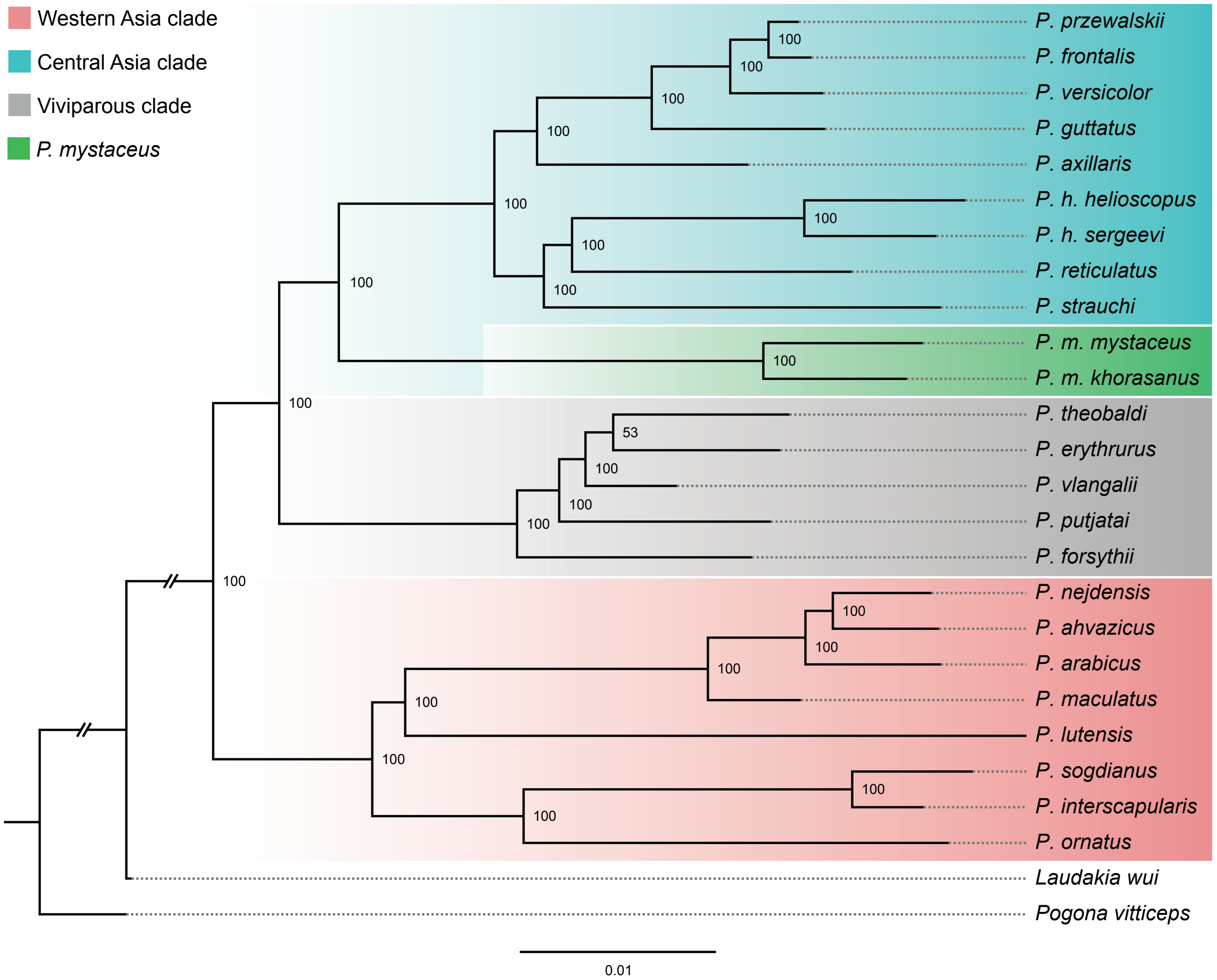


**Figure S1** Concatenation phylogeny inferred from protein-coding sequence. Maximum likelihood phylogeny reconstructed in IQ-TREE from the SNVs located in protein-coding regions.

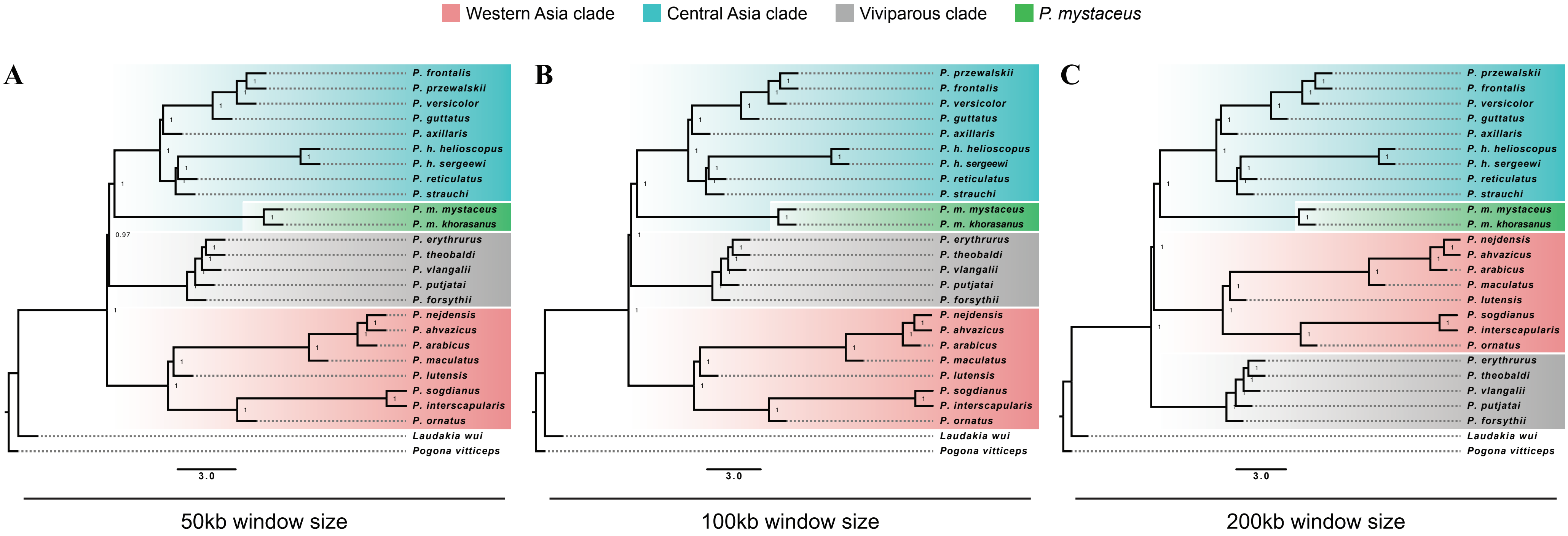


**Figure S2** Coalescent phylogenies reveal discordance among three major clades. Multispecies coalescent (MSC) species trees inferred with ASTRAL-III from local trees generated in fixed, nonoverlapping windows of 50kb (A), 100kb (B), and 200kb (C). Node labels indicate local posterior probabilities.

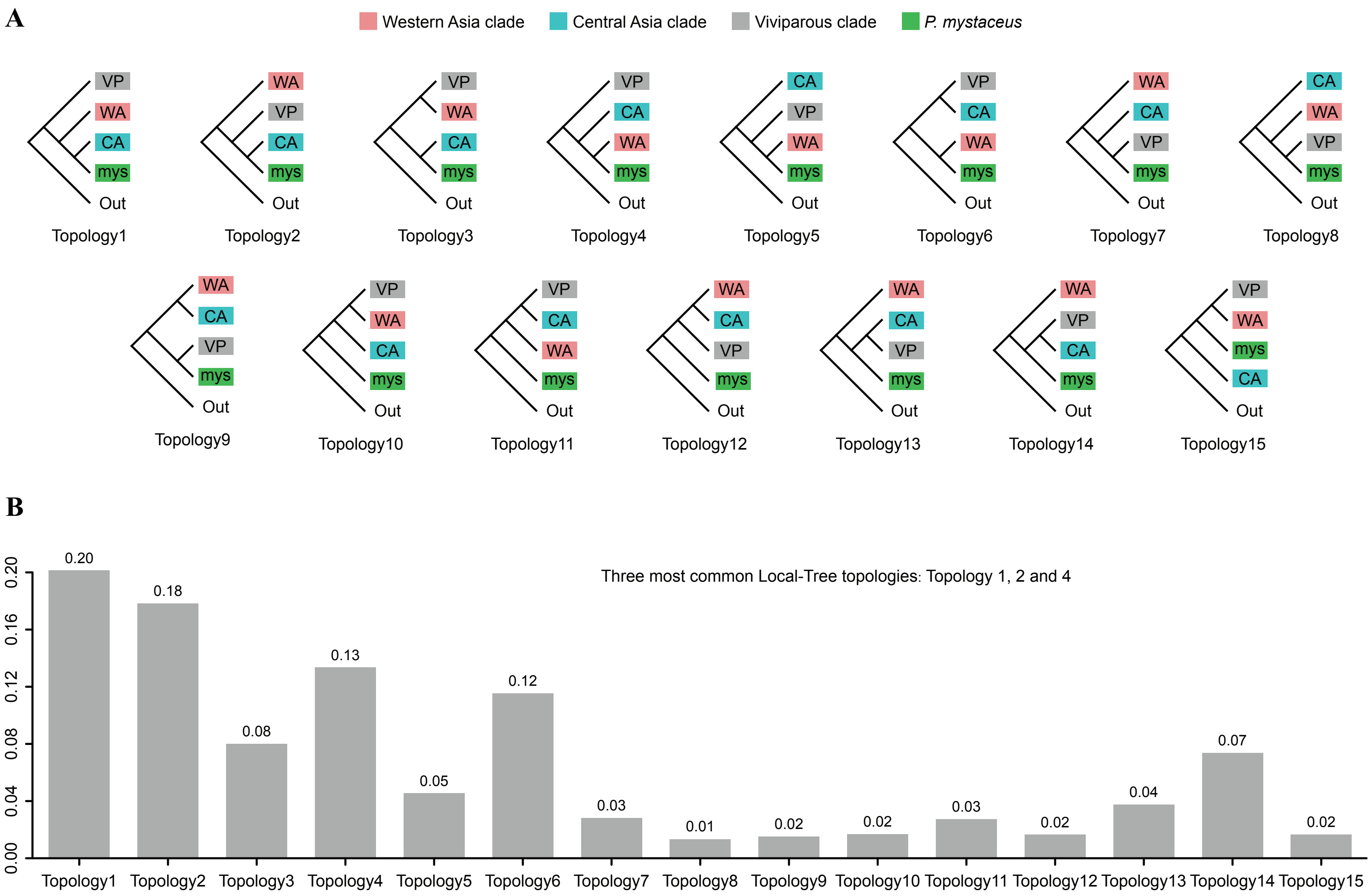


**Figure S3** Fifteen alternative inter-clade topologies retrieved from local tree analysis. (A) Classification of 8,662 local trees (fixed 200kb windows) into 15 distinct topologies relating the Western Asia, Central Asia, and viviparous clades plus *P. mystaceus*. (B) Genome-wide frequencies of each topology; the three most prevalent are Topo 1 (20.14%; congruent with the coalescent species tree), Topo 2 (17.83%; matching the concatenation tree), and Topo 4 (13.35%; placing *P. mystaceus* as sister to the Western Asia clade)

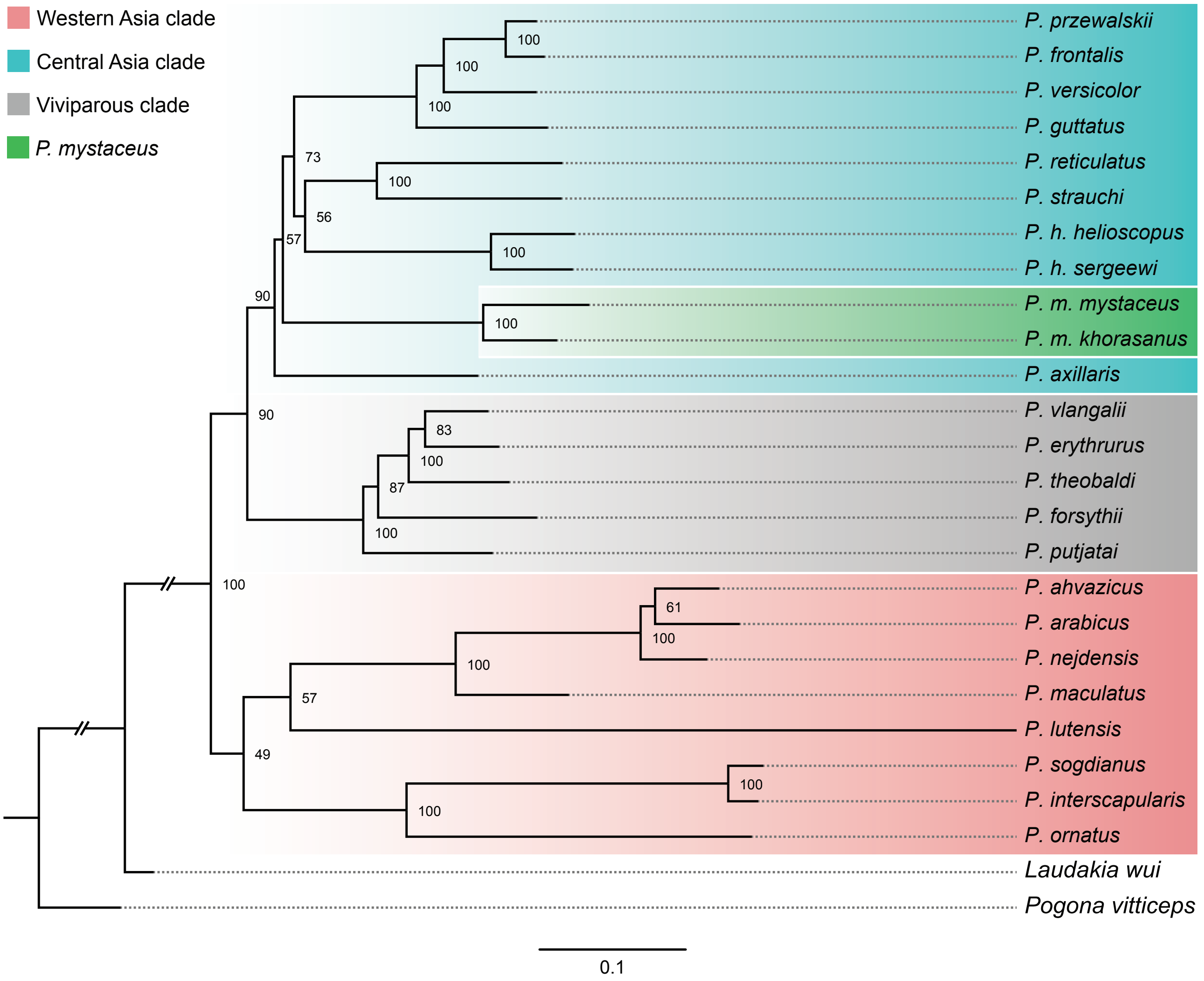


**Figure S4** Mitochondrial phylogeny inferred from 12 protein-coding genes. Maximum likelihood phylogeny reconstructed in PhyloSuite from the 12 mitochondrial protein-coding genes. IR denotes Iran; KA denotes Kazakhstan.

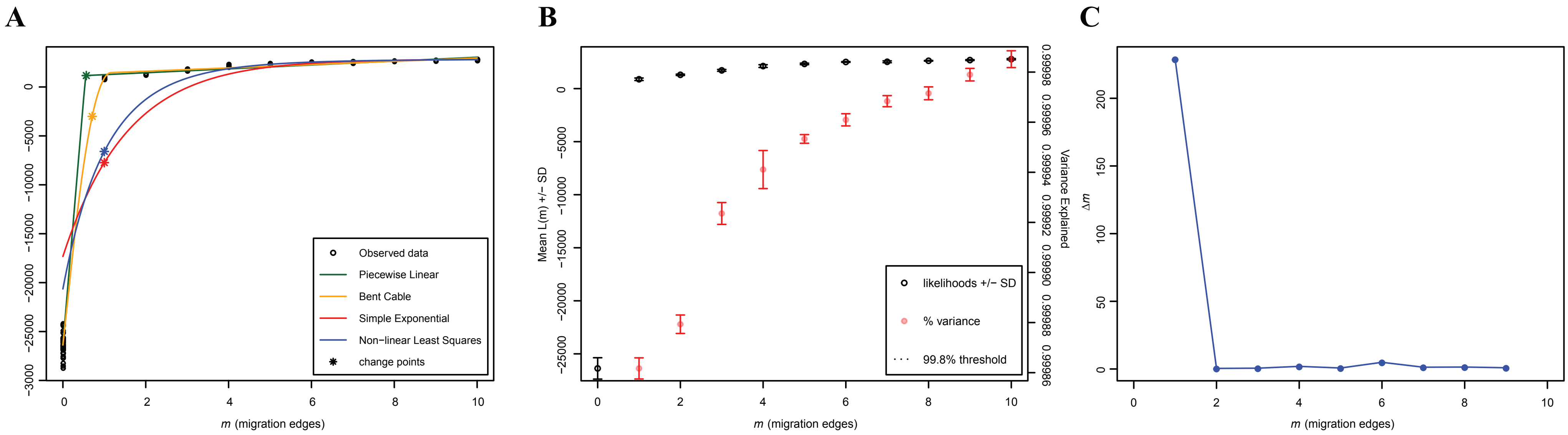


**Figure S5** One migration event identified as optimal by "OptM". The optimal number of migration edges (m = 1) in TreeMix was inferred with the R package "OptM" using (A–B) the "Evanno" method and (C) the "Linear" method.

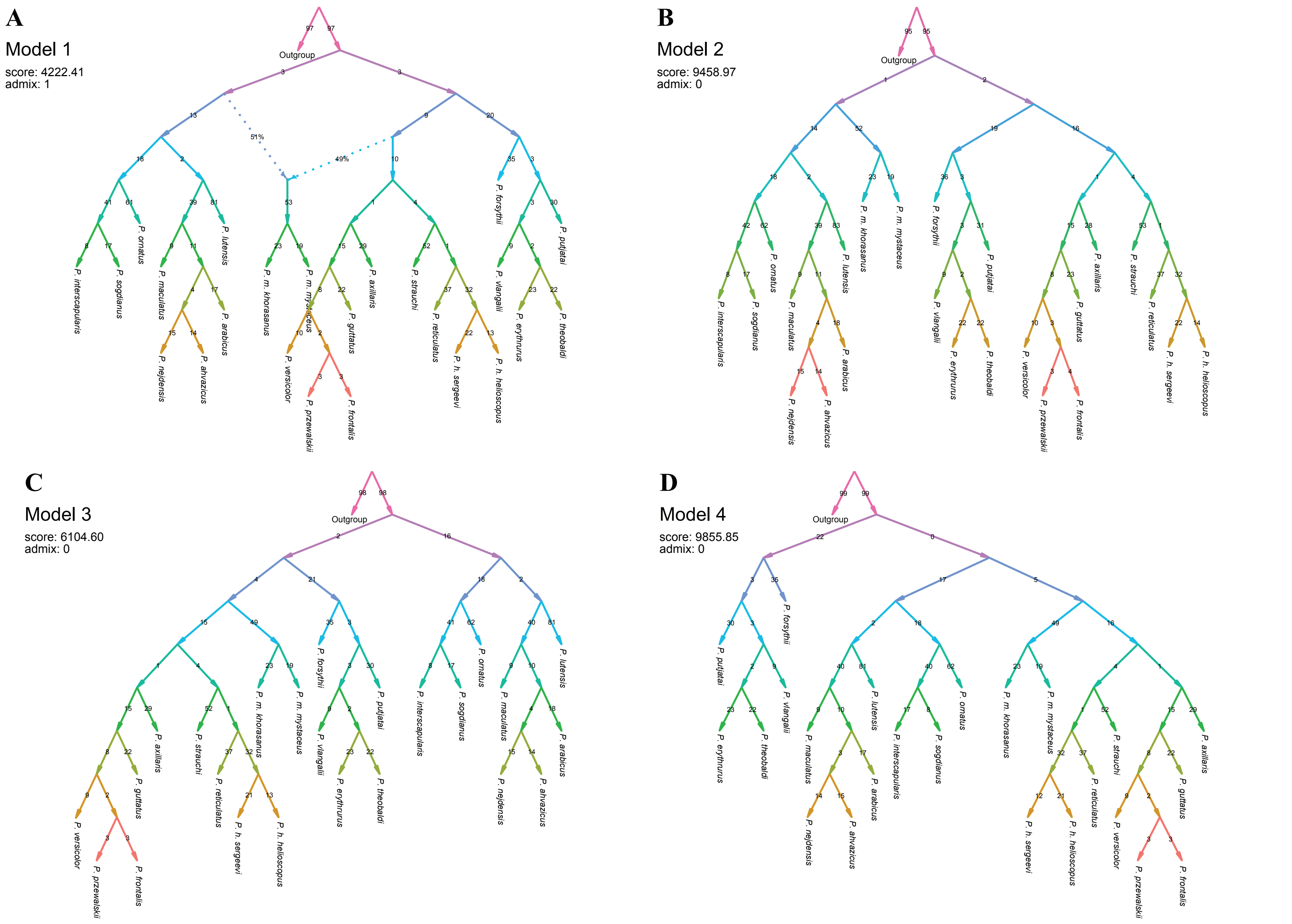


**Figure S6** Admixture graphs evaluated with ADMIXTOOLS 2. (A, Model 1) Best-fitting model including a single admixture event identified with "find_graphs" function (based on $f_3$-statistics computed from precomputed $f_2$-statistics). (B–D) Alternative no-hybridization models scored with: "qpGraph" function: (B, Model 2) concatenation topology; (C, Model 3) coalescent topology; (D, Model 4) an alternative topology placing *P. mystaceus* as sister to the Western Asia clade. Model 1 achieved the lowest score among the four models. IR denotes Iran, and KA denotes Kazakhstan.

**Table S1** Table of species used in this study

| Species | Sample ID | Latitude | Longitude | Accession |
|---|---|---|---|---|
| *P. maculatus* | ap01 | 30.74 | 55.94 | PRJCA045046 |
| *P. nejdensis* | ap02 | 29.54 | 35.42 | PRJCA045046 |
| *P. ahvazicus* | ap03 | 31.73 | 48.13 | PRJCA045046 |
| *P. ornatus* | ap04 | 33.91 | 59.39 | PRJCA045046 |
| *P. versicolor* | ap05 | 42.99 | 93.40 | PRJCA045046 |
| *P. frontalis* | ap06 | 40.21 | 111.12 | PRJCA045046 |
| *P. reticulatus* | ap07 | 40.26 | 63.86 | PRJCA045046 |
| *P. interscapularis* | ap08 | 41.75 | 64.04 | PRJCA045046 |
| *P. guttatus* | ap09 | 44.01 | 80.81 | PRJCA045046 |
| *P. axillaris* | ap10 | 42.86 | 89.19 | PRJCA045046 |
| *P. h. sergeevi* | ap11 | 40.71 | 66.53 | PRJCA045046 |
| *P. lutensis* | ap12 | 30.45 | 59.35 | PRJCA045046 |
| *P. h. helioscopus* | ap14 | 46.19 | 83.59 | PRJCA045046 |
| *P. przewalskii* | ap15 | 40.07 | 102.21 | PRJCA045046 |
| *P. arabicus* | ap16 | 18.51 | 53.14 | PRJCA045046 |
| *P. strauchi* | ap17 | 40.73 | 71.38 | PRJCA045046 |
| *P. m. khorasanus* | npm01 | 34.62 | 58.73 | PRJCA045046 |
| *P. m. mystaceus* | npm07 | 43.85 | 52.53 | PRJCA045046 |
| *P. sogdianus* | PhrSo | 37.42 | 67.51 | PRJCA045046 |
| *Laudakia wui* | LwThin | 25.23 | 95.14 | PRJCA045046 |
| *P. forsythii* | vp01 | 41.97 | 81.55 | PRJCA022059 |
| *P. vlangalii* | vp20 | 36.27 | 96.83 | PRJCA022059 |
| *P. erythrurus* | vp23 | 35.53 | 93.91 | PRJCA022059 |
| *P. theobaldi* | vp28 | 32.40 | 80.04 | PRJCA022059 |
| *P. putjatai* | GD069 | 36.04 | 101.49 | PRJCA022059 |
| *Pogona vitticeps* | PogVi | 28.10 | 144.47 | ERZ094017 |

**Table S2** Estimated divergence time of key nodes with geographic dispersal events

| Key Nodes | Posterior Means | 95% HPD |
|---|---|---|
| The common accoster of *Phrynocephalus* and *Laudakia* | 31.5656 Ma | 20.2318–43.5106 Ma |
| The crown node of *Phrynocephalus* | 17.6337 Ma | 12.6153–22.6546 Ma |
| The node of the Central Asia and viviparous clade | 15.8095 Ma | 11.3871–20.4736 Ma |
| The node of (*P. ornatus*, (*P. interscapularis*, *P. sogdianus*)) | 10.0734 Ma | 6.5633–14.4398 Ma |
| The node of ((*P. axillaris*, (*P. guttatus*, (*P. versicolor*, (*P. przewalskii*, *P. frontalis*)))), (*P. strauchi*, (*P. reticulatus*, (*P. h. helioscopus*, *P. h. sergeewi*)))) | 9.5467 Ma | 7.1993–11.4558 Ma |
| The node of (*P. maculatus*, (*P. arabicus*, (*P. nejdensis*, *P. ahvazicus*))) | 5.8651 Ma | 3.7561–8.1669 Ma |
| The node of (*P. guttatus*, (*P. versicolor*, (*P. przewalskii*, *P. frontalis*))) | 5.0045 Ma | 3.1660–6.8368 Ma |
| The node of (*P. ahvazicus*, *P. nejdensis*)) | 2.8430 Ma | 1.5673–4.2151 Ma |
| The node of (*P. m. mystaceus*, *P. m. khorasanus*) | 4.7456 Ma | 2.4484–7.2400 Ma |

**Table S3** Models and parameters of ancestral range estimation

| Models | LnL | numparams | d | e | j | AICc | AICc_wt |
|---|---|---|---|---|---|---|---|
| DEC | -38.85 | 2 | 0.011 | 1.00E-12 | 0 | 82.28 | 0.15 |
| DEC+J | -36.58 | 3 | 0.0057 | 1.00E-12 | 0.044 | 80.37 | 0.39 |
| DIVALIKE | -38.97 | 2 | 0.014 | 1.00E-12 | 0 | 82.52 | 0.13 |
| DIVALIKE+J | -36.8 | 3 | 0.0072 | 1.00E-12 | 0.035 | 80.81 | 0.31 |
| BAYAREALIKE | -48.8 | 2 | 0.017 | 0.063 | 0 | 102.2 | 7.20E-06 |
| BAYAREALIKE+J | -40.23 | 3 | 0.0041 | 1.00E-07 | 0.058 | 87.66 | 0.01 |

LnL: log-likelihood; numparams: number of free parameters; d: rate of range expansion (anagenetic dispersal along branches); e: rate of range reduction via local extinction/extirpation (along branches); j: relative weight of founder-event (jump) speciation at cladogenesis; AICc: Akaike information criterion corrected for small samples; AICc_wt: Akaike weights.
